# Multi-tissue transcriptome-wide association studies

**DOI:** 10.1101/2020.07.13.201111

**Authors:** Nastasiya F. Grinberg, Chris Wallace

## Abstract

Many genetic mutations affecting phenotypes are presumed to do so via altering gene expression in particular cells or tissues, but identifying the specific genes involved has been challenging. A transcriptome-wide association study (TWAS) attempts to identify disease associated genes by first learning a predictive model on an eQTL dataset and then imputing gene expression levels into a larger genome-wide association study (GWAS). Finally, associations between predicted gene expressions and GWAS phenotype are identified.

Here, we compared tree-based machine learning (ML) method of random forests (RF) with more widely used linear methods of lasso, ridge, and elastic net regression, for prediction of gene expression. We also developed a multi-task learning extension to RF which simultaneously makes use of information from multiple tissues (RF-MTL) and compared it to a multi-dataset version of lasso, the joint lasso, and to a single tissue RF. We found that for prediction of gene expression, RF, in general, outperformed linear approaches on our chosen eQTL dataset and that multi-tissue methods generally outperformed their single-tissue counterparts, with RF-MTL performing the best. Simulations showed that these benefits generally propagated to the next steps of the analysis, although highlighted that joint lasso had a tendency to erroneously identify genes in one tissue if there existed a disease signal for that gene in another.

We tested all four methods on type 1 diabetes (T1D) GWAS and expression data for several immune cells and found that 46 genes were identified by at least one method, though only 7 by all methods. Joint lasso discovered the most T1D-associated genes, including 15 unique to that method, but this may reflect its higher false positive rate due to “overborrowing” information across tissues. RF-MTL found more unique associated genes than RF for 3 out 5 tissues. Compared to lasso-based analysis, the RF gene list was more likely to relate to T1D in an analysis of independent data types. We conclude that RF, both single- and multi-task version, is competitive and, for some cell types, superior to linear models conventionally used in the TWAS studies.

**Author summary:** A transcriptome-wide association study (TWAS) is a way of integrating expression data and genome-wide association studies (GWAS), which allows for discovery of genes, rather than mutations, associated to traits of interest. In the TWAS framework, we first train predictive models on an eQTL dataset, then use these models to impute gene expression into a GWAS dataset. Finally, we look for significant associations between predicted gene expression and a GWAS trait. In this work, we compare non-linear method of random forests (RF) to linear models, customarily used in TWAS. Furthermore, we demonstrate that TWAS framework can naturally be extended to, and potentially benefit from, a multi-tissue setting, thereby taking advantage of the correlation between gene expression in different tissue types. We applied the RF, a selection of linear models, and the multi-tissue approaches to an eQTL dataset of monocytes and B cells and a large T1D GWAS. We found that RF outperform lasso in terms of predictive accuracy and the number of differentially expressed genes found, and that multi-dataset version of lasso discovered the most T1D-associated genes. Analysis of the gene lists produced for each method in independent data types (excluding genetic association data) showed all related to T1D, but that the RF methods ranked T1D higher in their lists than the linear methods. We conclude that RF is a useful addition to the TWAS tool box.

## Introduction

Genome-wide association studies (GWAS) have been hugely successful over the last decade, transforming genetic association testing into a reproducible science [1] and identifying tens of thousands of variants associated with more than a thousand traits [2]. However, lack of interpretability remains a criticism of GWAS [3]—most disease-associated variants lie in regulatory regions [4, 5] but have not yet been convincingly linked to the genes they regulate. It has been noted that eQTLs are over-represented among trait-associated SNPs uncovered by GWAS [6, 7]. This has motivated development of different methods to link GWAS variants to genes by integrating GWAS and eQTL datasets [8–10], and one promising approach, referred to as transcriptome-wide association study (TWAS), is to use an eQTL dataset to learn rules with which to impute gene expression in GWAS samples. Predicted gene expressions can then be used in place of genotypes within the standard GWAS framework, enabling gene-based instead of variant-based, case-control comparisons [11].

Previously proposed approaches for learning the imputation rules are based on regularized linear models [11–14], polygenic risk scores [11] and using the top SNP to predict expression levels [12]. However, the machine learning literature has shown that alternative approaches such as random forests (RF), which allow naturally for non-linear and non-additive effects, can produce more accurate predictions in model organisms [15, 16]. We set out to explore whether using RF could also lead to better gene expression predictions in humans and, if so, whether that could be translated into a more powerful TWAS.

We also sought to take advantage of the fact that expression levels of a given gene in different cell types can be correlated by considering expression values across multiple cell types simultaneously in a multi-task framework. This has been shown to improve multi-trait predictions in yeast [16] and in applications to real and simulated data in marker-assisted selection for several related traits [17–19] or populations [20]. Multi-trait approaches have also been used to analyse eQTL datasets [21, 22]. We adapted standard RF for this purpose and compared it to the joint lasso of Dondelinger and Mukherjee [23], as well as to the linear and RF models trained on data from single tissue only.

We compared linear and RF models for single-tissue and multi-tissue learning and studied how their performance translated into TWAS using simulated data and a real data example. For this, we chose to consider eQTL data from multiple immune cell types (B cells and monocytes [24, 25]) for a large type 1 diabetes (T1D) study [26].

## Results

### Random forests allow improved predictions of gene expression in single tissues

We began by comparing the predictive power of different methods of eQTL prediction in a microarray gene expression dataset of monocytes and B cells from 430 individuals [24, 25] (Table 1). For each probe, SNP markers within 1 Mbp of that probe (*cis*-SNPs) were used to train a predictive model for each cell type. Only probes which have at least one cell type with a nominally associated *cis*-SNP (*p*-value < 10^−7^; see) were considered (4,288 probes resulting in 21,440 probe-cell regressions; see Methods for more details)

**Table 1.** Summary of eQTL datasets used in this study. Expression data of Fairfax *et al* [24, 25] includes B cells and monocytes, unactivated and activated—response to interferon-γ (IFN) and lipopolysaccharide after 2 (LPS2) and 24 (LPS24) hours.

| Dataset | Cell type | Samples | SNPs | Probes |
| --- | --- | --- | --- | --- |
| Fairfax *et al* | CD14<sup>+</sup> | 413 | 588,141 | 47,230 |
|  | CD14<sup>+</sup> LPS2 | 260 |  |  |
|  | CD14<sup>+</sup> LPS24 | 321 |  |  |
|  | CD14<sup>+</sup> IFN | 366 |  |  |
|  | B cell | 284 |  | 47,231 |

Models were trained on a training set and evaluated on a test set, comprising roughly 70% and 30% of the data, respectively. Predictive accuracy was assessed by calculating *R*^2^ on the test set. In order to avoid information leaking in the MTL set-up, described later, all samples from the same individual were designated to either the training or the test set.

We compared performance of RF [27] to three regularised regressions: lasso [28], ridge [29] and elastic net [30], motivated by their popularity in the literature. Lasso and ridge regressions differ by their use of an *L*^1^ or *L*^2^ penalty parameter, respectively, with elastic net being a mixture of the two.

Amongst the linear methods ridge regression strictly underperformed compared to lasso and elastic net which performed similarly to each other, with lasso slightly preferred (Fig S2 (a)), suggesting that eQTL prediction benefits from sparsity introduced by the elastic net and lasso regression. Moreover, once sparsity is introduced, varying the mixing parameter hardly affected performance of elastic net (Fig S2 (b)). We therefore dropped ridge regression and elastic net from further analysis.

RF outperforms lasso in the overwhelming majority of regressions with mean advantage (see Methods) of RF over lasso of 5.9%, compared to 3.5% of mean advantage of lasso over RF (Figure 1). Moreover, for 1,927 out of 11,814 probe-cell pairs with any signal, RF beat lasso by more than 10%. A cluster of points near the origin and along the *x*-axis in the second quadrant of the RF-lasso graph demonstrates that RF detected signal in some of the regression problems for which lasso only sees noise (Fig 1).

**Fig 1.**
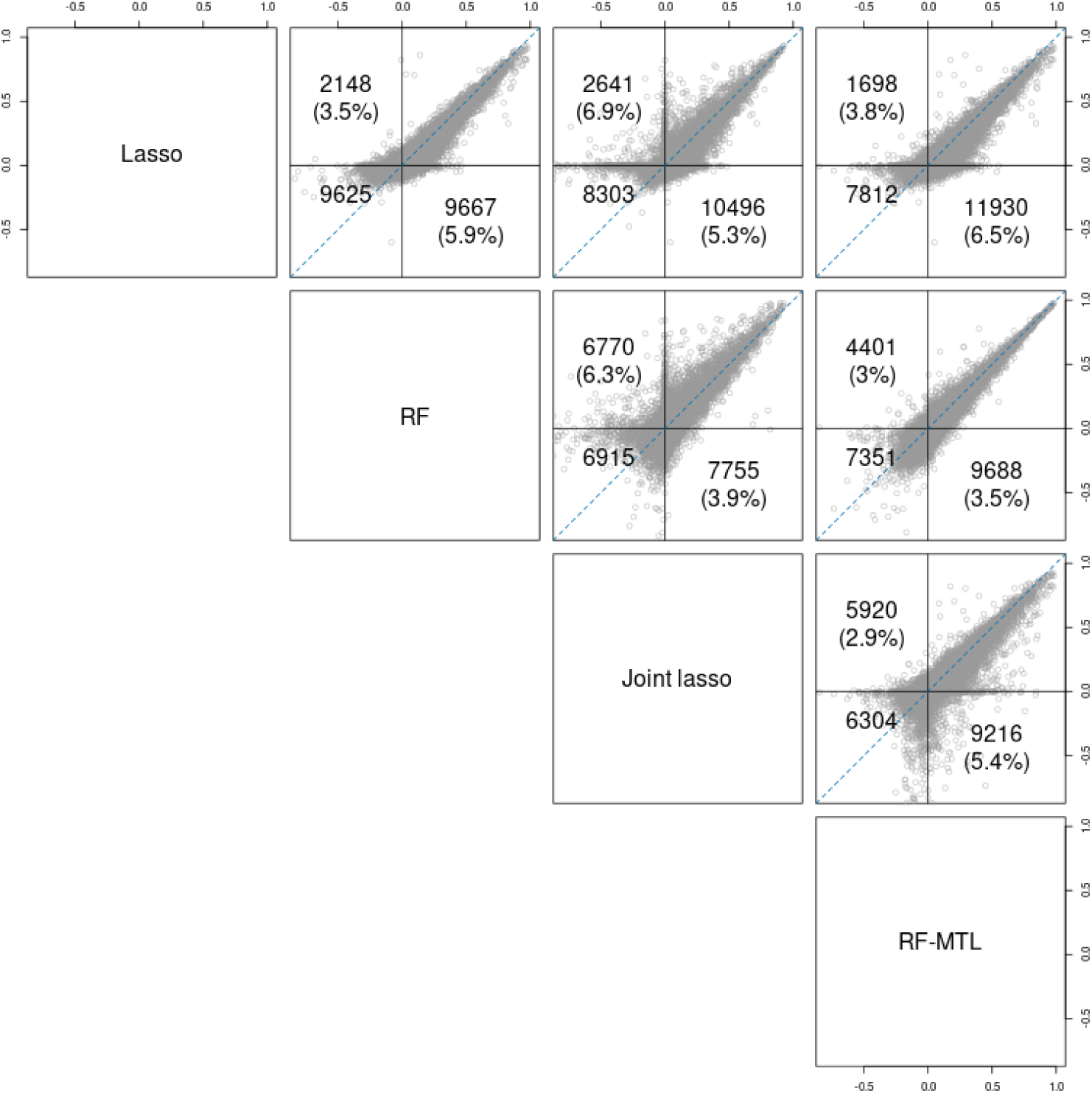
Pairwise comparison of performance of the MTL and STL expression prediction methods—*R*^2^ on a test set. Each point represents a probe-cell pair. Points above the blue line show increased performance for the method to the left of each plot, while points below the blue line show increased performance for the method underneath the plot. The three numbers represent, clockwise: points with positive *R*^2^ above *x* = *y* line for the *x*-axis method, points with positive *R*^2^ below the line for the *y*-axis method, points with negative *R*^2^ for both methods. Numbers in brackets represent the corresponding advantage of one method over the other, in terms of *R*^2^ (for this calculation negative *R*^2^ are taken to be 0). For example, comparing lasso and RF, lasso outperformed RF in 2,148 regressions with an advantage of 3.5%, while RF outperformed lasso in 9,667 with an advantage of 5.9%, and for 9,625 probe-cell pairs neither method achieved a positive *R*^2^.

### Combining information from multiple cell types using multi-task learning

Multi-task learning (MTL) leverages similarities between targets of several regression problems by learning these targets simultaneously [31, 32]. Moreover, it is known that many eQTLs are active across multiple cell types [33]. We, therefore, examined whether MTL could improve performance of both RF and lasso using data from multiple cell types sampled from the same individuals. For this, we trained RF on all five cell types simultaneously and tested performance of an MTL version of lasso—joint lasso of Dondelinger and Mukherjee [23]. We compared the two to each other and to the reference models fitted on individual tissue types (single-task learning; STL). We considered the same 4,288 probes for which at least one cell type has a nominally associated *cis*-SNP (*p* < 10^−7^) *p*-value, resulting in the same number of regressions (each able to predict expression for five cell types).

To implement an MTL version of RF (we refer to it as RF-MTL), for each probe we pooled expression levels across all cell types into one large regression problem, adding a categorical variable indicating tissue of origin as an extra variable. Note that this variable was included as a candidate for splitting at each split and for each tree. The test set consisted of the same 30% of the samples used for testing in the single-task setting.

For RF, the pooled approach above should cater for situations when the underlying sub-datasets have a varying degree of similarity. Pooling completely homogeneous (or even identical) datasets, should not adversely affect performance as the tissue id variable, although available as a splitting variable at every split, does not have to be used if it does not help reduce residual variance for a given tree. Strong differences between sub-groups, on the other hand, should be handled by the use of the tissue id variable at various splits, effectively separating samples from heterogeneous sub-groups. This, of course, stems from the assumption that similarities/dissimilarities between different sub-groups are reflected in similarities/dissimilarities of their respective distributions over features. Joint lasso handles multiple datasets simultaneously by estimating different regression coefficients for different tissues while encouraging coefficients of similar tissues to be closer. This is done by introducing an extra regularisation term penalising difference between coefficients of different sub-groups (*L*^1^ or *L*^2^ penalty) depending on how similar these sub-groups are with respect to a given dissimilarity measure.

Comparing joint lasso to standard lasso (Fig 1) we see that the former outperforms the latter in the absolute majority of cases. However, joint lasso significantly underperforms in a handful of cases, against lasso as well as RF and RF-MTL. RF-MTL and RF are relatively evenly matched, although RF-MTL performs slightly better in more regressions. RF-MTL outperforms joint lasso substantially more often than the other way around (in 9,161 and 5,918 regressions, respectively) and tends to have a larger advantage (5.4% compared to 2.9%). Overall, RF-MTL, on average, is the most accurate predictive model for our eQTL dataset. Additionally, only one regression has to be fitted to cater for all cell types instead of one per cell type.

### Simulation-based comparison of learning methods for TWAS

To complete a TWAS analysis, we need to not simply detect associated genes, but also remove cases where the associations result from distinct eQTL and GWAS variants in LD. For this, we add as a second stage a filtering step. If causal variants are shared, we expect the pattern of association for the two traits will mirror each other, such that coefficients will be proportional across SNPs in the region [34]. We therefore use a proportional colocalisation test [35] to filter out TWAS-significant associations that are not consistent with proportionality. We define a *TWAS-significant* association as a cell-probe-method triplet for which predicted expression has a significant fold change (as assessed by the FDR-adjusted Cochran-Armitage test *p*-value).

To assess the performance of the four methods as part of the complete two-stage TWAS procedure we simulated GWAS-trait and gene expression data for five cell types under several genetic causal scenarios. One of the cell types was designated “test”, while the other four cell types were designated as “background”, and either all or none of the “background” cell types shared causal variants with either “test” cell type and/or the GWAS trait. For each replicate, we trained all four predictive models above, predicted expression for the “test” cell type in the GWAS dataset, and tested association between GWAS-trait and predicted expression. For identified associations, we also tested for proportionality of genetic effects on GWAS and expression in the associated cell type [35]. This test is expected to preferentially filter out significant TWAS results that result from an eQTL variant distinct from, but in LD with, a GWAS causal variant. We evaluated the methods based on the proportion of TWAS-significant calls in different scenarios before and after filtering.

Generally, when colocalised GWAS and eQTL signals were simulated, multi-trait methods outperformed single-trait methods when eQTL variants were shared between the test and background expression traits, and single-trait methods performed slightly better when there was no sharing, though the difference was more pronounced in the former versus the latter (Fig. 2, top panels). However, the situation was very different when background expression traits shared a variant with the GWAS but the test expression trait did not. Here, we might expect an increase in false positives due to occasional LD between GWAS-trait variants and test-expression-trait variants, possibly explaining the higher false positive rate for unfiltered RF-MTL compared to RF (0.14 and 0.10, respectively). However, joint lasso performed particularly poorly in this scenario, with a false positive rate of 0.58 compared to 0.040 for single-task lasso. Testing proportionality was successful at preferentially filtering out false positives, reducing type 1 error rates to at or below their nominal value with the exception of the joint lasso case, where the false positive rate was only reduced to 0.37. Proportionality filtering also removed between 7.5% and 10.5% of true positives, fairly evenly across methods.

**Fig 2.**
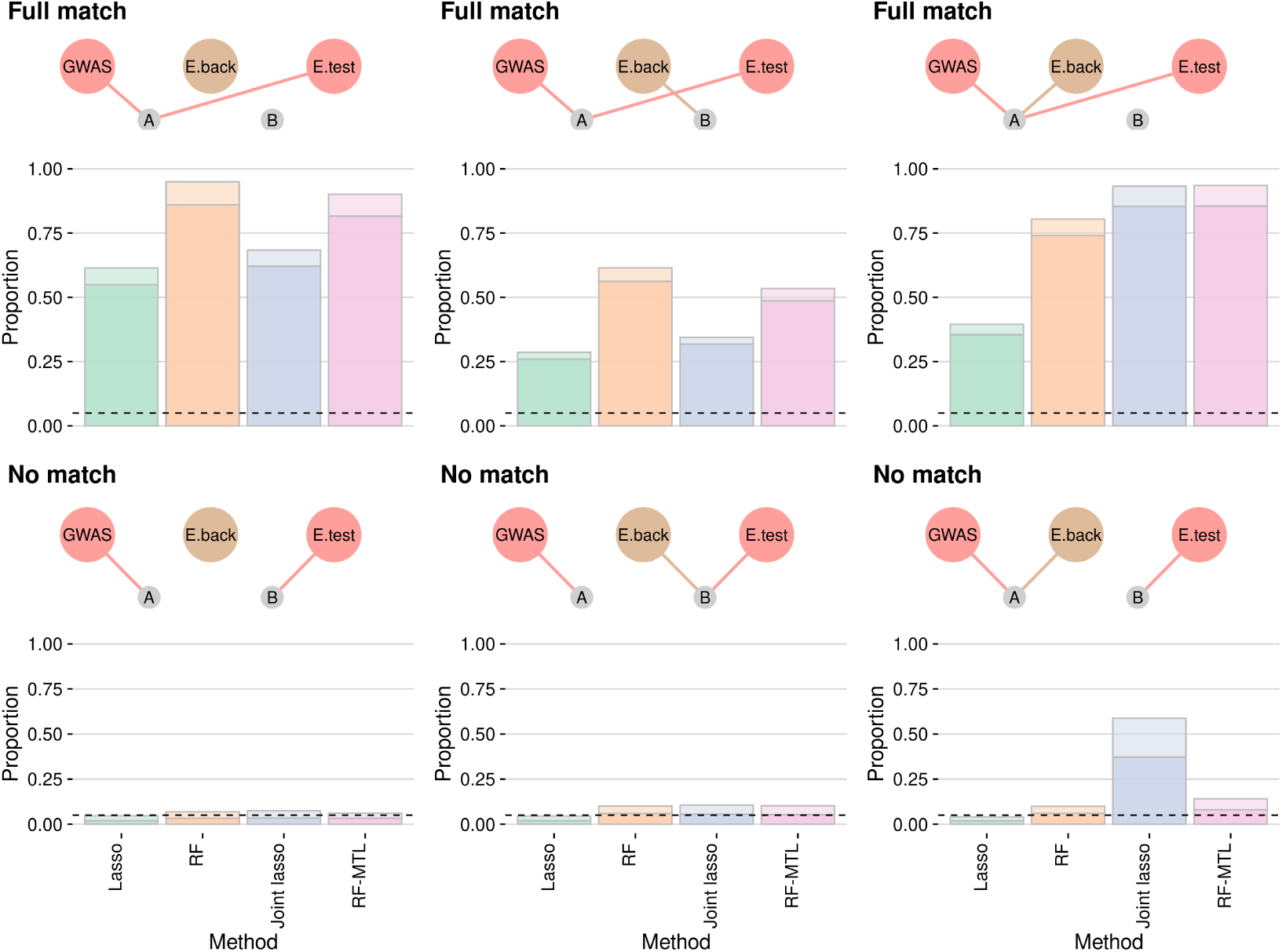
Power of different methods to detect TWAS association. In the top row, the GWAS and test eQTL traits share causal variant A, while the causal variant for the four background eQTL traits varies (left-right) from none, to B to A. The bottom row is the same, except the GWAS and eQTL-test causal variants are different. The total shaded column height is the proportion of TWAS tests that pass *p* < 0.05, with lighter shading used to indicate the proportion of tests which would be filtered out proportionality testing at *p* < 0.05.The horizontal dotted line is at *y* = 0.05, the proportion of false positives expected in a well controlled testing procedure in the bottom row.

Overall, this suggests that the benefits of RF-MTL over RF, and of RF over lasso for prediction transfer to TWAS. On the other hand, they warn that joint lasso may have a high false positive rate if interpreted in a tissue specific manner. A more detailed comparison of single-task RF and lasso showed that the effects of regularisation on lasso caused systematic over-estimation of the causal effect of expression on the GWAS trait with lasso (S5 Fig).

### 46 genes show predicted differential expression in T1D

To compare performance of the predictive methods in a real-world dataset, we retrained the models on the whole eQTL data (as opposed to 70% training set) and used them to impute (predict) gene expression into a large T1D GWAS cohort [26] (5,913 cases and 7,341 controls). We tested for a difference in mean predicted gene expression between cases and controls using a Cochran-Armitage test, stratified by the two studies the data is comprised of (see Table S1).

Overall, 62 distinct TWAS-significant genes were identified by at least one of the four methods with joint lasso identifying the most (see Table 2, column 4). Filtering for proportionality left 46 distinct genes (Table 2, Figure 3). We call TWAS-significant hits passing the proportionality filter *SP-hits* (significant and proportional). There is a substantial overlap between the four methods but each also identified unique hits not discovered by the others (Figures 4 and S6). RF finds an equal or greater number of unique SP-hits than lasso in all but one cell type. Likewise, RF-MTL finds at least as many or more unique SP-hits than single-tissue RF in three out of five tissue types. Joint lasso identifies the most TWAS-significant and SP genes for each cell type but the heatmap of results 5 shows that joint lasso genes tend to be significant in three and more tissue types. Indeed, multiple full vertical lines designate instances when a gene is significant in all the cell types (see Discussion).

**Table 2.** Table of results of the TWAS analysis. Non-null regressions (N) refer to the expression prediction models taken through to the GWAS imputation state, i.e. lasso and joint lasso models which identify no useful SNPs, and hence offer only constant predictions, are dropped. TWAS-significant hits refer to predicted gene expressions passing the Cochran-Armitage test (5% with Benjamini-Hochberg adjustment) for differential expression in T1D. Finally, last column is the number of TWAS-significant hits passing the proportionality filter (at 5%)—SP-hits.

| *Method* | *Cell* | *N* | *TWAS-significant (unique)* | *SP-hits (unique)* |
| --- | --- | --- | --- | --- |
| Lasso | BCELL | 1155 | 25 (18) | 10 (8) |
| RF | BCELL | 4103 | 17 (10) | 8 (6) |
| Joint lasso | BCELL | 3886 | 44 (36) | 22 (19) |
| RF-MTL | BCELL | 4103 | 17 (11) | 6 (5) |
| Lasso | CD14<sup>+</sup> | 1962 | 14 (11) | 8 (6) |
| RF | CD14<sup>+</sup> | 4103 | 15 (12) | 8 (6) |
| Joint lasso | CD14<sup>+</sup> | 3485 | 32 (26) | 19 (15) |
| RF-MTL | CD14<sup>+</sup> | 4103 | 20 (15) | 10 (7) |
| Lasso | IFN | 1919 | 14 (10) | 5 (4) |
| RF | IFN | 4103 | 30 (24) | 13 (11) |
| Joint lasso | IFN | 3494 | 40 (32) | 22 (18) |
| RF-MTL | IFN | 4103 | 23 (18) | 10 (9) |
| Lasso | LPS2 | 1317 | 10 (8) | 5 (3) |
| RF | LPS2 | 4103 | 11 (10) | 5 (4) |
| Joint lasso | LPS2 | 3762 | 33 (29) | 17 (15) |
| RF-MTL | LPS2 | 4103 | 21 (16) | 11 (9) |
| Lasso | LPS24 | 1525 | 16 (13) | 4 (3) |
| RF | LPS24 | 4103 | 13 (11) | 6 (5) |
| Joint lasso | LPS24 | 3645 | 35 (31) | 21 (19) |
| RF-MTL | LPS24 | 4103 | 19 (15) | 10 (9) |
| *Total (unique)* |  |  | 449 (62) | 220 (46) |

**Fig 3.**
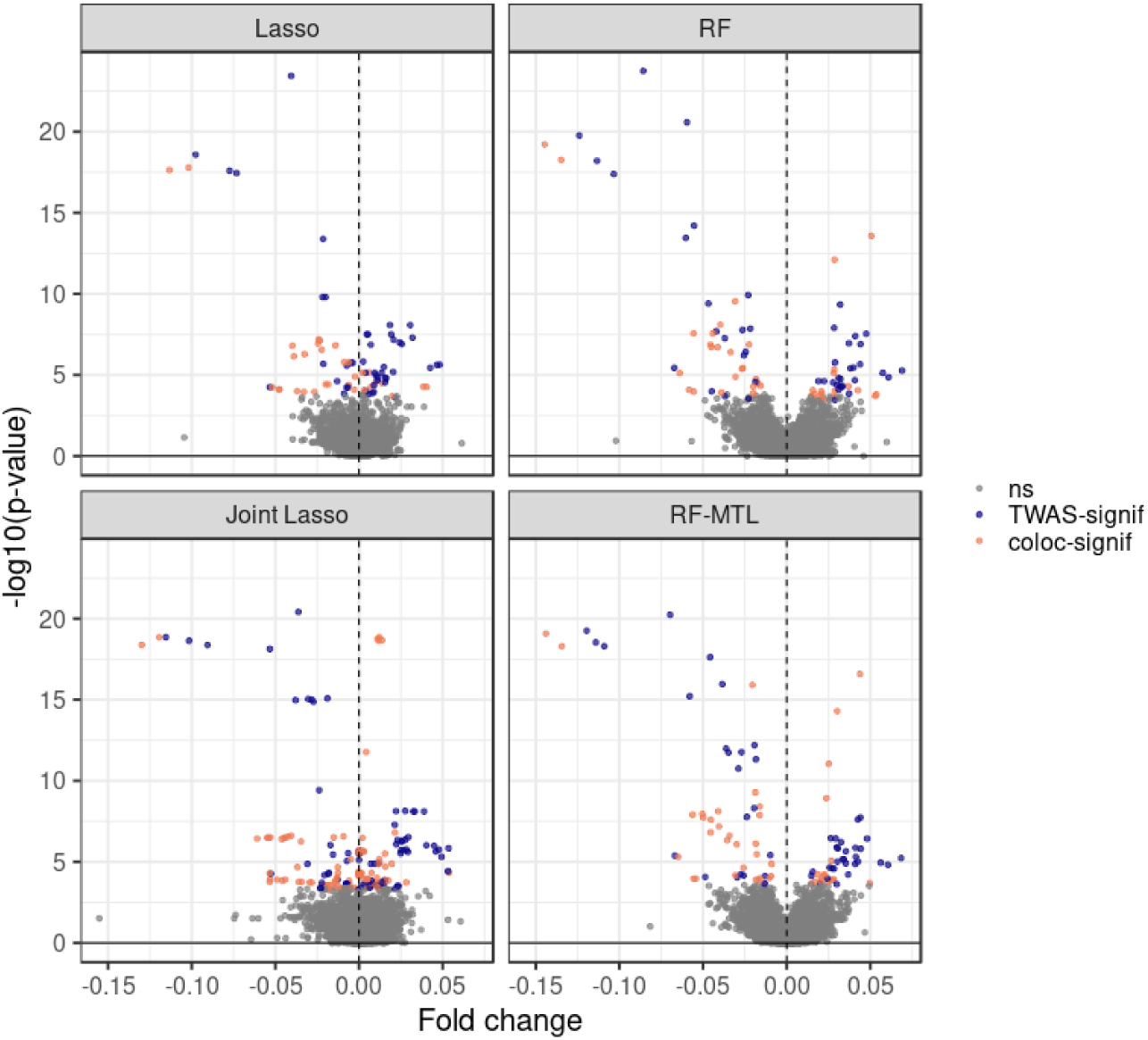
Volcano plots for testing association between the gene expression predicted by the four methods and the T1D status. Grey points are not TWAS-significant, blue points are TWAS-but not passing proportionality test, and orange points are both TWAS- and proportionality-significant (SP-hits).

**Fig 4.**
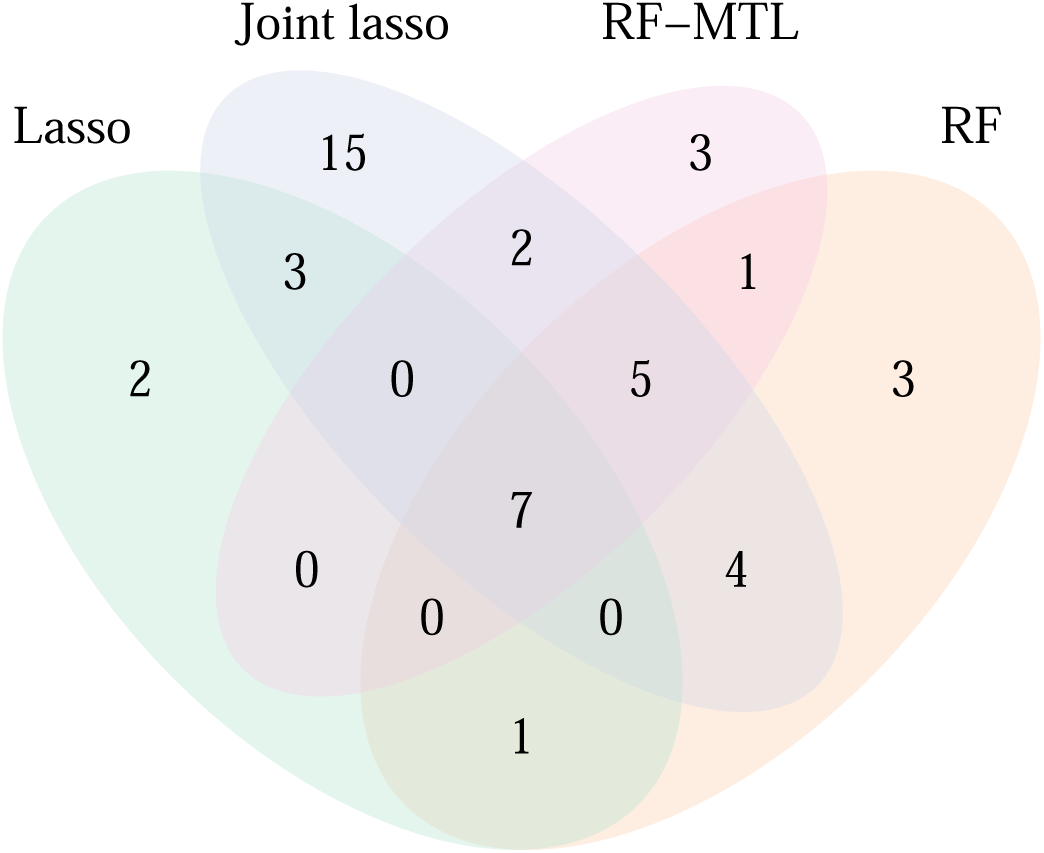
Unique TWAS-significant hits passing proportinality filtering, by method: lasso (13), RF (21), joint lasso (36), and RF-MTL (18).

**Fig 5.**
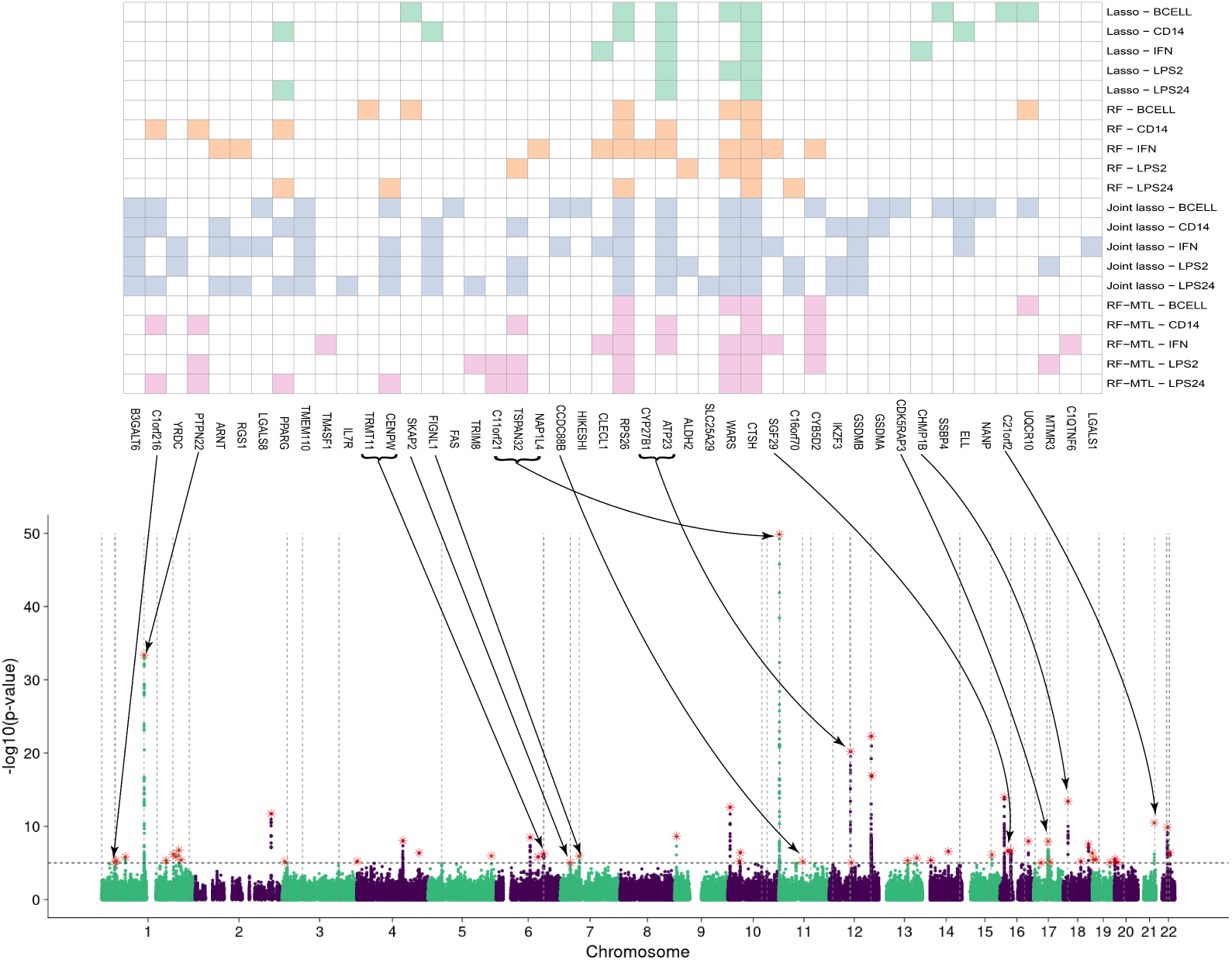
A heatmap of genes identified by the four methods after proportionality filtering (top), integrated with a manhattan plot of T1D GWAS. Arrows point to GWAS peaks (red stars) in the vicinity of which (1 mb either way) a gene (or several genes, grouped by a bracket) lies. Vertical dotted lines indicate positions of genes; horizontal dotted line is at *y* = 5, corresponding to a GWAS significant level; green and purple colours in the manhattan plot designate alternating chromosomes. Note that the genes in the heatmap are ordered according to their positions, so for any two genes (or groups of genes) an arrow form a leftmost one would point to a peak left of the peak pointed at by the rightmost gene. Any intersection between the arrows is due to the fact that they might point to peaks of vastly different heights.

As the complete list of true T1D genes is not known, we decided to compare the results from the different methods by passing the gene list to the Target Validation web analysis platform (https://www.targetvalidation.org/) and searching for associated diseases, excluding genetic association data from the data types included to avoid circular reasoning. We ranked the diseases listed according to their relevance *p*-value, and found that the RF-based gene lists ranked more obviously T1D-related diseases higher than lasso-based gene lists (table S2). Indeed, the term “type I diabetes mellitus” was the second ranked for RF and the third ranked for RF-MTL, but only the 19th for lasso (19th) and 45th for joint lasso (45th), supporting that RF-based TWAS was identifying disease-relevant genes identified by methods independent from genetic association data.

## Discussion

The aim of TWAS is to associate genes and diseases. Although association can be thought necessary for causation, it is not sufficient [36]. We use colocalisation analysis to determine whether, for a TWAS-significant gene, the same genetic signal underlies the eQTL and a trait-association, or whether two (or more) distinct signals exist in some linkage disequilibrium (LD). We do this via testing for proportionality of SNP regression coefficients for the two traits in question [35]. This alternative framing of the null hypothesis differs from the more widely known enumeration method for colocalisation [37] (where the null hypothesis is no association for either trait) and is a more natural way to approach this question once a joint association has been found. Our approach is thus related to the two-stage HEIDI/SMR approach proposed by Zhu *et al* [9]. Colocalisation validation was also used in [10, 13]. However, recently other methods of validating/fine-mapping TWAS signals have been proposed—Mancuso *et al*. [38], for example, extend probabilistic SNP-level fine-mapping approaches to create credible sets of genes which explain a given TWAS signal with a given probability.

We note that associated genes filtered for lack of proportionality would be expected to be differentially expressed in healthy individuals at different risks of disease (those who carry greater or lesser burdens of disease-predisposing variants). Thus, we might expect them to also be differentially expressed between cases and controls in a hypothetical study in which expression is measured directly. Therefore, we suggest such genes might be considered as biomarkers rather than red herrings. Even SP-hits can be validated only through practical lab-based experiments.

The current ubiquity of linear methods in eQTL studies reflects both the speed and flexibility of these methods, but also the prevailing dogma that gene expression is influenced additively over variants and alleles at those variants. This expectation reflects the lack of evidence from human studies directly targeting epistatic effects [39–41]. However, this lack of evidence could also reflect a lack of power [42]. While exploiting RF was not unreservedly a more powerful method for TWAS, the fact the RF predictions were generally better than those from lasso suggests that non-additive effects make an important contribution in gene expression. Such non-linearity has been detected in detailed molecular studies of individual genes [43], and in large scale studies of model organisms [44]. It also motivates wider development and adoption of methods that can exploit non-additivity where it exists, even in samples insufficiently large for non-additivity to be robustly detected.

It is important to understand the reasons behind differences in performance of the four methods, both in terms of predictive accuracy and the number of TWAS-significant hits discovered. Both tree-based methods outperformed their linear counterparts on average, with the RF-MTL being the most accurate overall. Clearly, whilst the lasso methods are competitive, RF-based methods successfully exploit the supposed non-linear relationships in the data. For T1D, however, this predictive advantage did not translate into more TWAS-significant hits consistently across different tissue types. The reason for may lie in the fundamental differences in the properties of the two models. Lasso (and so, joint lasso) produces biased solutions (unlike standard linear regression) with the resulting coefficients biased towards zero, accepting this cost in order to generate predictions with lower variance. Random forest, on the other hand, produces a low-bias model but higher variance predictions (see Figures S3 and S4). As a consequence, even lasso predictions resulting in very small fold changes can lead to TWAS-significant hits through incorporating few (sometimes just one) but important SNPs in predictive models (i.e. highly biased but low variance predictions). This results can be seen most clearly comparing the shape of the volcano plots (Figure 3), where the expected dip in the middle is not evident in lasso. Overall lower variance of RF-MTL predictions but similar size of predicted fold change, as compared to RF, might also explain why RF-MTL does better in the TWAS framework.

Multi-tissue methods demonstrated their applicability to TWAS both in terms of accuracy of models constructed on the eQTL dataset and the number of unique TWAS-significant genes and SP-genes associated to TID identified. Indeed, Hu *et al*. [22] found that their multi-tissue method UTMOST outperformed single-tissue elastic net (PrediXcan [11]) in both stages of the TWAS framework. Like joint lasso, the UTMOST predictive model is a type of regularised regression with several penalty terms in addition to the standard least squares loss. The two penalties used in UTMOST are: *L*^1^ for effect sizes within each tissue for variable selection and effect size shrinkage, and *L*^2^ grouped lasso penalty for effect sizes across tissues to encourage cross-tissue eQTLs. RF-MTL, on the other hand, uses expression data from different tissues in a flexible non-parametric manner, exploiting similarities where they exist.

The effects of regulatory variation have been shown to vary between cell types [24], and cell type specific chromatin accessibility has been used to associate multiple immune cell types to autoimmune disease GWAS [45]. Hence, for a given disease, it is important not only to identify potential genes of interest but also the relevant tissue(s). Simulations showed that the two multi-tissue methods we studied tend to “overborrow” information across tissues, i.e. find significant hits for tissues without one if there is a real signal in another tissue. This was mostly a problem suffered by joint lasso and, to a much smaller extent, by RF-MTL. It is harder to identify this behaviour in real data. However, the number of TWAS-significant hits identified by joint lasso in our T1D data and the fact that it was much more likely to find signal in 3 or more tissues for a given gene than the other methods, suggests similar behaviour. Moreover, calculated standard deviation of predicted fold change for different cell types for each probe (for lasso methods, for probes with at least three cell types with non-null predictions) reveal that joint lasso has the least variation in fold change predictions between different tissue types (see Figure S7). Hence, whilst outperforming single-tissue lasso on average in terms of prediction accuracy, joint lasso seems to suffer from lower prediction specificity and, as a result, a higher rate of false positive TWAS-hits in the TWAS framework.

In this study, we demonstrated applicability of non-linear and multi-tissue methods in the TWAS framework. Both real data and simulation studies showed, in particular, that RF is at least as competitive and, for some tissue types, superior to lasso. Similarly, RF-MTL is superior to RF for some tissue combinations. Our results highlight the potential to exploit multiple tissue-eQTL studies in TWAS but we expect this to be most useful when tissues are closely related, so that information may be legitimately borrowed between tissues.

## Methods

### Measure of accuracy

Throughout the paper we use *R*^2^ (*R*-squared) as a measure of predictive accuracy of different models. For a predictive model *f, R*^2^ is informally known as the ‘proportion of the variance explained’ by *f* and is defined as:

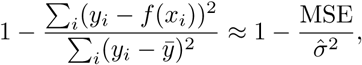

where *f* (*x*_*i*_) is prediction at point *x*_*i*_, 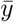 is sample mean of outcome *y*, 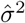 is *y*’s sample variance and MSE is mean square error. Note that the above fraction is a measure of how well *f* does compared to the ‘base’ constant model 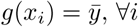. One would expect a ‘good’ model to have small MSE compared to 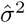, and hence larger *R*^2^. Conversely, a ‘bad’ model will have a larger MSE and smaller *R*^2^, with a truly hopeless model performing en par with a constant mean predictive function. Note also that, whilst the phrase ‘proportion of variance explained’ would entail a value of *R*^2^ in the interval [0, 1], in reality the definition above does not put any such restriction on *R*^2^. Indeed, a heavily overfitting model, or that trained and tested on data coming from vastly different distributions, can produce large negative *R*^2^ values.

For two methods, *m*_1_ and *m*_2_, trained and validated on the same datasets with respective *R*-squared, 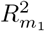 and 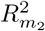, we say that *m*_1_ *has an advantage over m*_2_ if 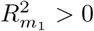 and 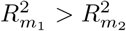. This advantage is quantified by 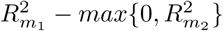. *Average advantage* of *m*_1_ over *m*_2_ is calculated over a set of regression problems to which both methods are applied and *m*_1_ has an advantage over *m*_2_. In case of Pearson correlation being used as an accuracy method, average advantage is defined analogously.

### Selection of probes for study

For efficiency, the first step of our analysis is to filter probes with no genetic predictability. Even though standard univariate eQTL association analysis, by virtue of its linearity, does not show the full picture of relationships between SNPs and expression, it is fast and can help us to gauge the strength of genetic signal for each probe. We, therefore, only keep those probes that have at least one *cis*-SNP with a *p*-value of less than 10^−7^ for at least one cell type. Additionally, we excluded the HLA region (chr6:20mbp-40mbp). Probe positions, originally on build 38 (GRCh38), were lifted over to build 18 (NCBI Build 36.1) to match the genotypic data. Some probes could not be matched and were discarded. Hence, out of the original 47,231 probes 25,005 survived the liftovers, and only further 4,288 passed the *p*-value thresholding and were retained for analysis.

### eQTL prediction

All expression values used in the STL models (elastic net, RF) were standardised to have mean 0 and variance 1, individually for each cell type. For the MTL framework (joint lasso, RF-MTL), for each eligible probe, we centered the expression values to have mean 0 (but did not standardise them) for each cell type individually.

### Elastic net

Lasso and ridge regression do not require explicit tuning: the complexity parameter *λ* is chosen via internal cross-validation when the model is fit. Elastic net, being a mixture of the two, has an additional parameter *α* ∈ [0, 1] with *α* = 1 corresponding to full lasso and *α* = 0 to full ridge. Usually, the mixture parameter *α* is also tuned via cross-validation, but often a fixed value is chosen, e.g. Gamazon *et al* [11] simply use *α* = 0.5. However, in our experience predictive accuracy of elastic net is a monotonic function of *α* (when complexity parameter *λ* is tuned through cross-validation separately for each value of *α*, as opposed to doing a two-dimensional grid search). Usually the best performance is exhibited by one of the extremes, 0 (ridge) or 1 (lasso), with the performance for 0 ≤ *α* ≤ 1 being very similar to that of the model with the winning *α* parameter. We demonstrate this here for the eQTL dataset of Fairfax *et al* (using all five tissue types). We assessed performance of the elastic net model for a range of the mixture parameter *α* evenly spaced between 0 and 1. Figure S2 depicts violin and boxplots of *R*^2^ on a test set for each *α* for each probe/cell pair.

### Joint lasso

We opted for the *L*^2^ fusion version of the joint lasso as it requires less tuning compared to the *L*^1^ fusion, and the original paper [23] reported a similar performance for both. We tuned the *L*^2^ joint lasso for the fusion parameter *γ* via external 5-fold cross-validation and for the general penalty parameter *λ* via in-built internal 10-fold cross-validation (i.e. within each fold of the *γ*-tuning fold, lasso would tune for *λ* via another cross-validation routine). For any probe and two tissues *i* and *j* we set group specific penalty *τ*_*ij*_ to *ρ*_*ij*_*/max*_*k*≠*l*_ {*ρ*_*kl*_}, where *ρ*_*ij*_ is the correlation between expression of *i* and *j* in the Fairfax dataset. However, in [23], authors remark that in practice using non-constant (unity) *τ* ‘s didn’t improve predictive performance of joint lasso. The joint lasso was implemented using the fuser package.

### Random forest

RF requires relatively little tuning: the optimal number of trees is determined by assessing out of bag error as the forest is grown (we grew 500 trees which was sufficient for convergence) whilst it has been suggested that regulating depth of the trees (via minimum number of observations in terminal nodes) has limited benefits [46, 47]. We incline to agree, as tuning RF simultaneously both for tree depth and the number of variables considered for splitting at each split made little difference to the accuracy of the resulting model (results not shown). We thus used the default parameter values: minimum number of observations in terminal notes at 5 (resulting in deep trees), and the number of random variables considered at each split at a 1*/*3 of all SNPs (parameters `min.node.size` and `mtry`, respectively). We used the `ranger` function in the `ranger R` package to fit RF.

### RF-MTL

To implement multi-trait prediction in RF, we simply combined (stacked) expression matrices for the five tissue types into one tall matrix. Then, each individual could have up to five associated sample points. Genotypic matrices were similarly stacked and an id variable indicating which tissue/dataset each point came from was added. This variable was available for splitting at each iteration of the RF algorithm (`always.split.variables = “id”` in the `ranger` function). This way the size of the training data was increased and the underlying structure could be taken advantage of, or ignored, depending on its presence.

### Gene expression imputation

The four predictive models (lasso, RF, joint lasso, RF-MTL) are trained on the full eQTL datasets but only using those *cis*-SNPs that also feature in the corresponding GWAS T1D dataset. For some probes no SNPs are shared between the two datasets, so out of the initial 4,288 probes we are left with 4,103. We then test for association between these imputed expression levels and the disease status of the individuals in the GWAS dataset, to see which probes/genes are differentially expressed—we use the Cochran-Armitage [48] test (with Mantel adjustment to account for stratification; see Table S1). To account for multiple testing, the resulting *p*-values are adjusted using the Benjamini-Hochberg [49] method (separately for each method and cell type). Note that for the two lasso methods the total number of fitted models, as opposed to just the non-null ones, were used for the *p*-values adjustment. This was done to avoid giving lasso and joint lasso an unfair advantage over the two forest models. TWAS-significant hits are those probe/cell pairs with adjusted *p*-value< 0.05.

### Proportionality filtering

To reduce the degrees of freedom of the test, proportionality testing works by first finding principal components (PCs) of the genotype matrix accounting for the majority of the variation (usually 80%), regressing the two traits on these PCs, and comparing the corresponding coefficients. Finally, a null hypothesis that the two sets of coefficients are proportional (there is a colocalisation) is tested. To reduce the number of PCs used, we only used SNPs with *p*-values< 10^−4^ and all the SNPs in their LD pockets (*r*^2^ > 0.2 with selected SNPs). We then use the PCs accounting for at least 80% of the variation, or the first 6 PCs, whichever number is the smallest. 13 out of 224 TWAS-significant probe/cell pairs (6 probes) did not have enough SNPs with sufficiently small *p*-values and were dropped.

### Simulations

We sampled independently 400 pairs of haplotypes from the 1000 Genomes EUR subset to generate genotype data, and sampled causal variants independently from amongst the SNPs according to the scenarios described in Figure 2.

6 quantitative traits were simulated as Gaussian variables with variance 1 and mean ∑_*i*_ *β*_*i*_*G*_*i*_ where *i* indexes causal variants, *β*_*i*_ is the effect size and *G*_*i*_ the genotype. To avoid too many simulations with small beta and non-significant effects, *β*_*i*_ was sampled as the maximum of 5 Gaussians with variance 0.04. The first trait was assigned as a GWAS trait, the second the expression trait to be tested via TWAS, and the remainder as additional “background” expression traits. Each expression trait was regressed against all SNPs, and the simulation retained if the minimum *p*-value over all SNPs and expression traits was less than 10^−7^. TWAS was conducted with each of the 4 methods described above, and the *p*-value retained. We also ran proportional filtering, as described above, and stored its *p*-value.

We assessed TWAS performance according to the proportion of simulations that gave a TWAS *p*-value < 0.05, before and after filtering.

### Software

All analysis was done in `R` using `glmnet` for lasso and elastic net, `ranger` for RF and RF-MTL, and `fuser` and bespoke helper functions https://github.com/stas-g/fuser_helper for the joint lasso. coloc package was used for the post-hoc colocalisation analysis. All simulation code is available from https://github.com/chr1swallace/twas-sims.

## Supporting information

significant proportional hits

## Acknowledgements

This work was funded by the Wellcome Trust (WT107881) and the MRC (MC UP 1302/5).

## Supplementary Tables and Figures

**Fig S1.**
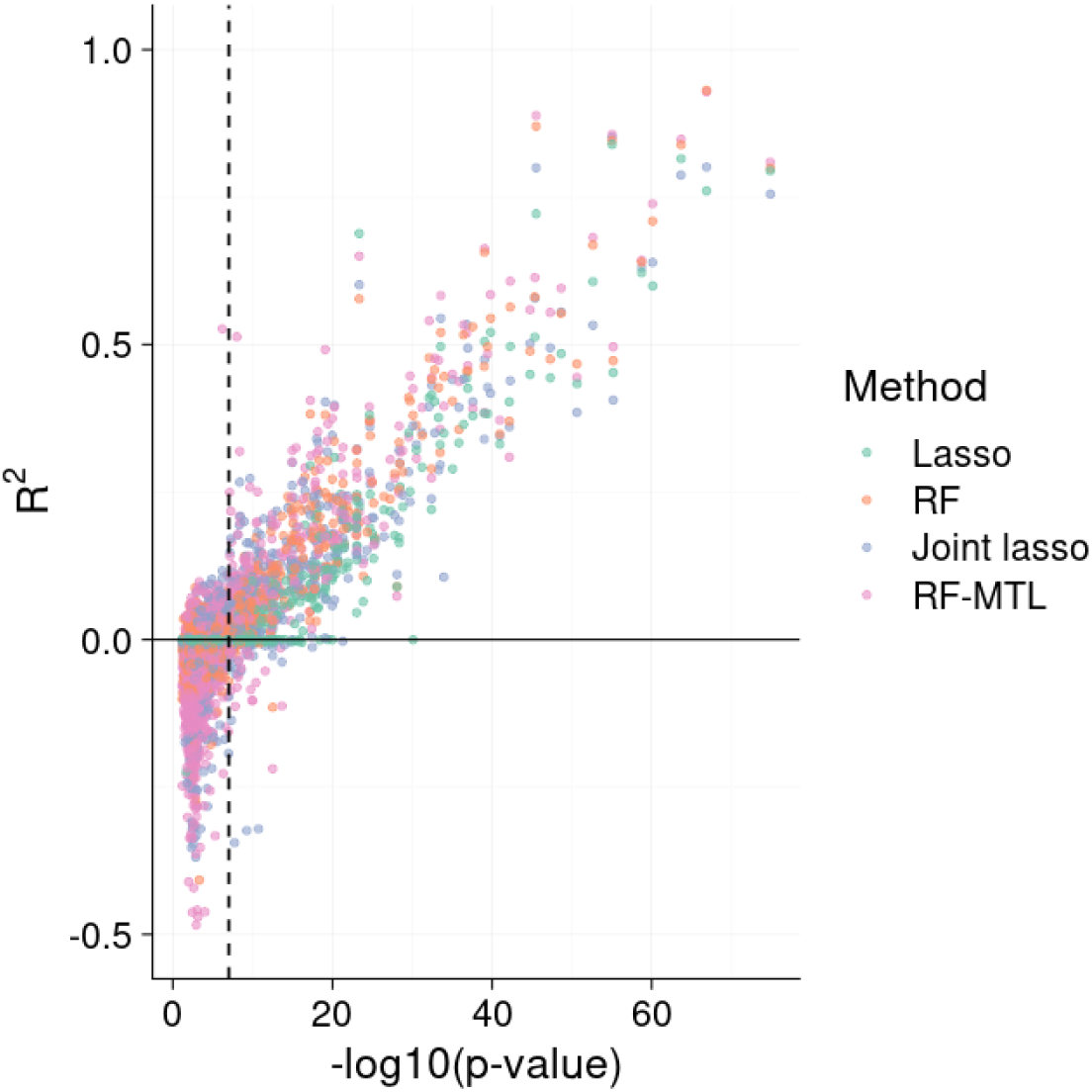
Performance of the four expression prediction methods, as assessed by *R*^2^ on a test set, plotted against the minimum *p*-value of the eligible (cis) SNPs for each probe/cell pair on chromosome 22 (3040 regressions for each method). The vertical dashed line is at *x* = 7sed (i.e. minimum *p*-value = 10^−7^).

**Table S1.**
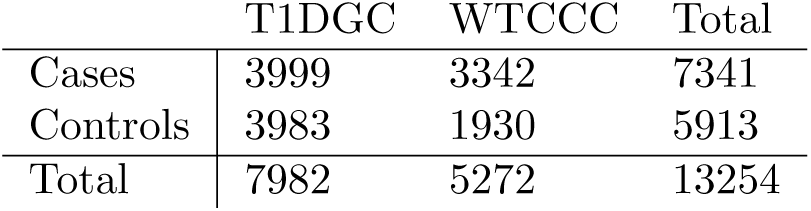
T1D data of Barrett *et al* [26] comprising Wellcome Trust Case Control Consortium (WTCCC) [50] and Type 1 Diabetes Genetics Consortium (T1DGC) samples.

|  | T1DGC | WTCCC | Total |
| --- | --- | --- | --- |
| Cases | 3999 | 3342 | 7341 |
| Controls | 3983 | 1930 | 5913 |
| Total | 7982 | 5272 | 13254 |

**Fig S2.**
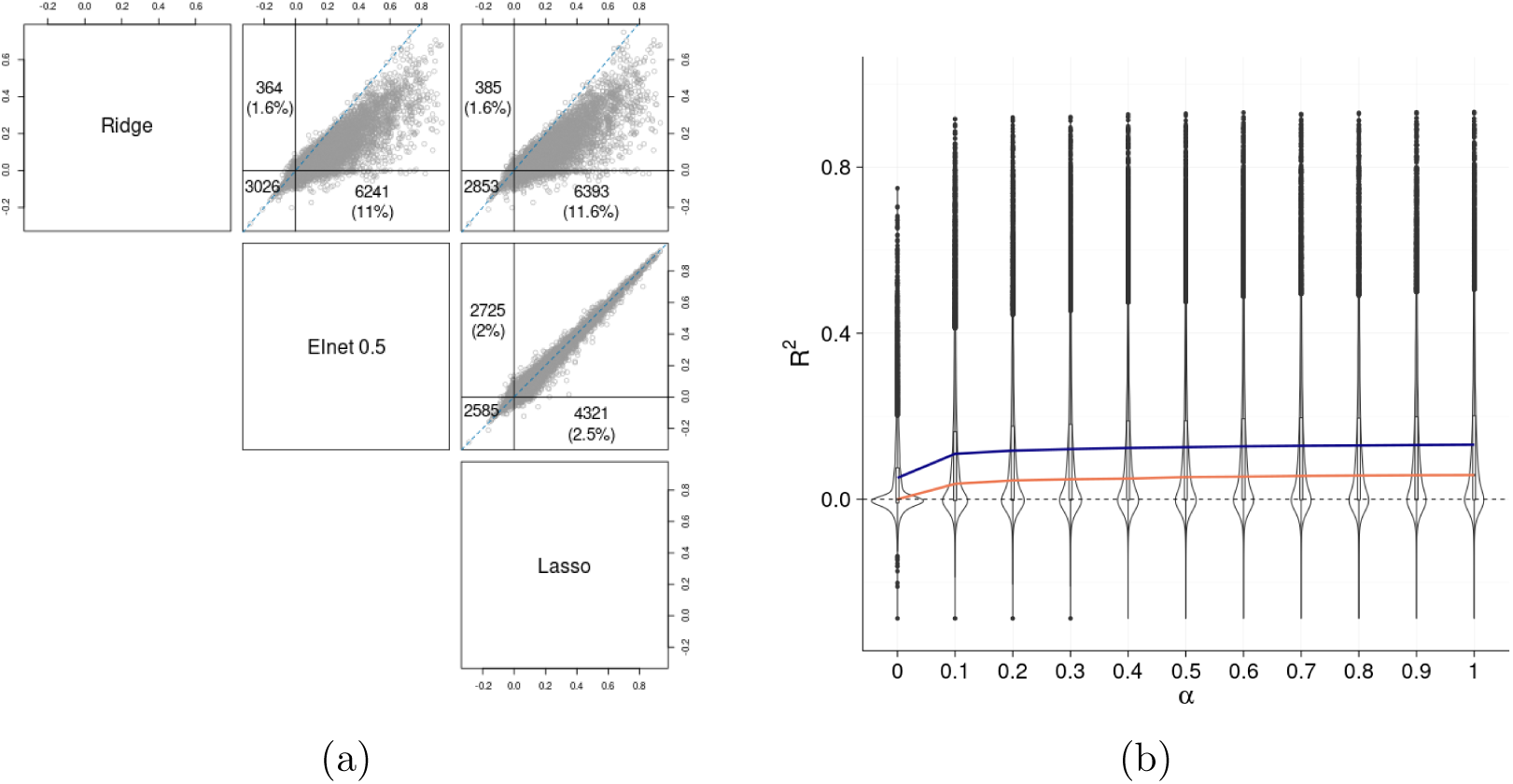
(a) Pairwise comparison of performance (*R*^2^ on a 30% test set) of elastic net for *α* = 0, 0.5, 1. Each point represents a probe-cell pair. Points above the red line show increased performance for the method to the left of each plot, while points below the red line show increased performance for the method underneath the plot. The three numbers represent, clockwise, starting top left: points with positive *R*^2^ for the *x*-axis method above the *x* = *y* line, points with positive *R*^2^ for the *y*-axis method below the line, points with negative *R*^2^ for both methods; average advantage in brackets. (b) Performance of elastic net for varying values of *α*, evenly spaced between 0 and 1, on the eQTL dataset of Fairfax *et al* (*R*^2^ on a 30% test set). Note that the values 0 and 1 correspond to the ridge regression and lasso, accordingly. Each violin plot, with the embedded boxplot, aggregates all regressions for a given *α*. The purple and orange lines are mean and median values of *R*^2^, respectively.

**Fig S3.**
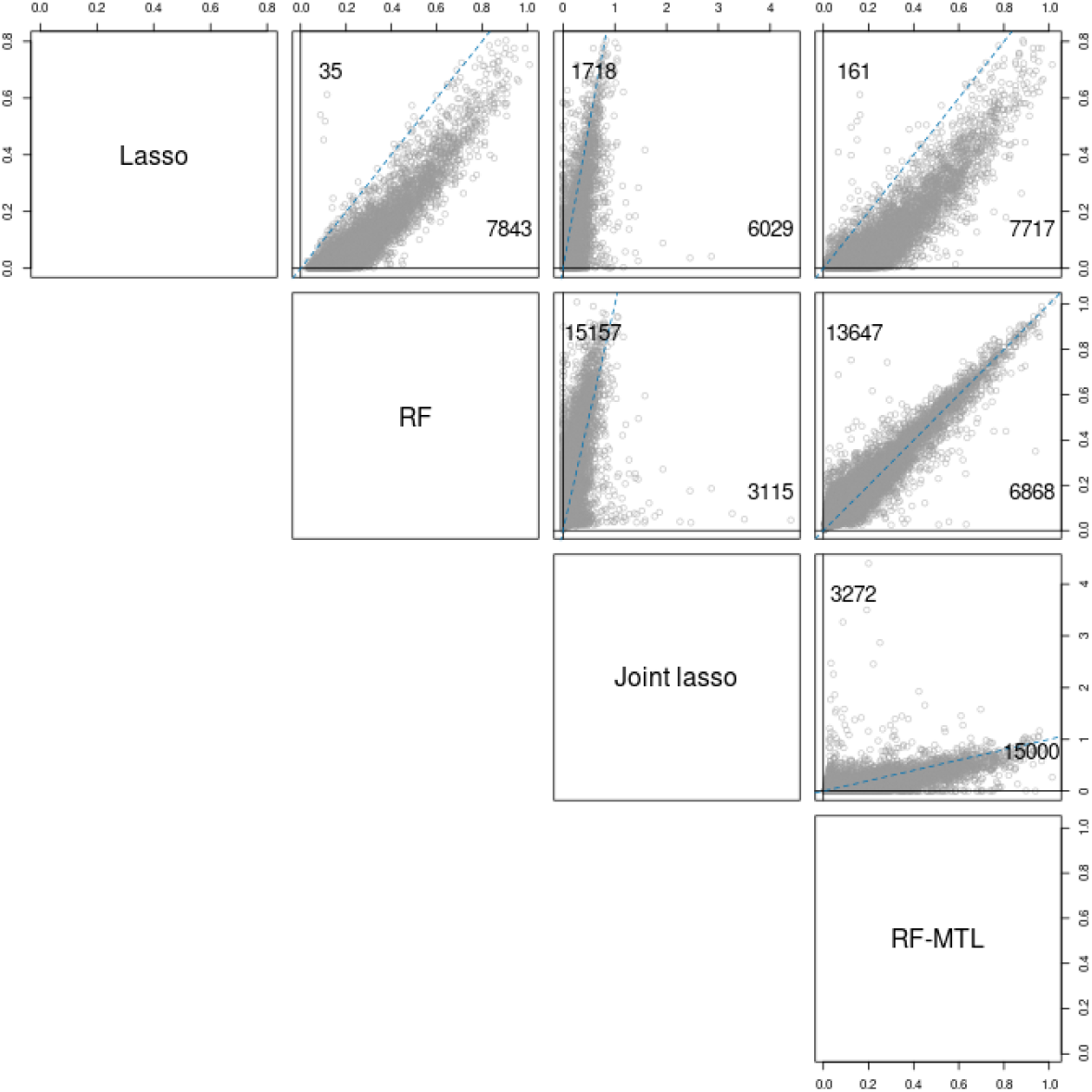
Pairwise comparison of variance of imputed expression values for the four methods. The blue dashed line is the *x* = *y* line. Numbers above and below the line correspond to the number of regressions for which the *y*-axis method has larger variance for the imputed predictions than the *x*-axis method and vice versa, respectively.

**Fig S4.**
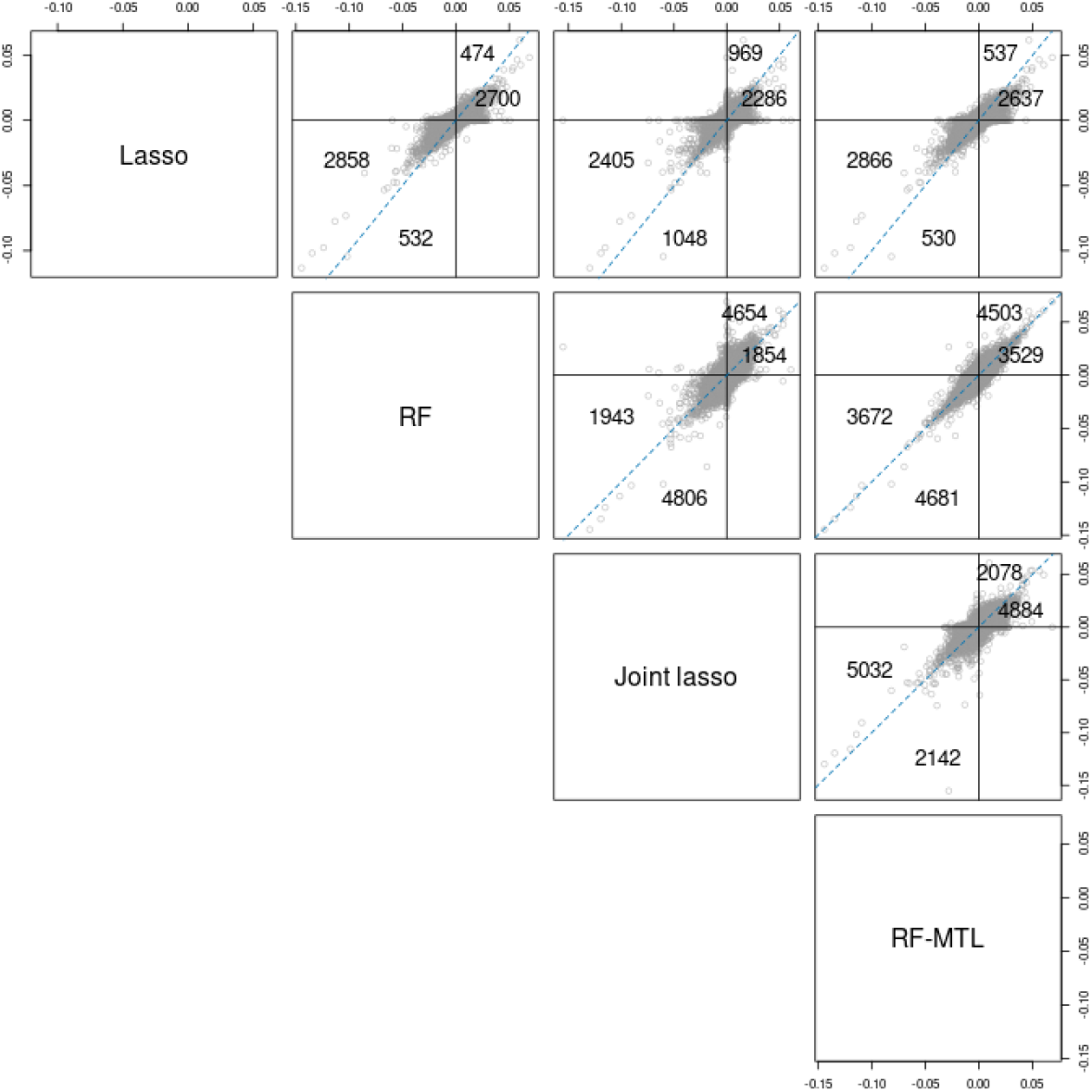
Pairwise comparison of predicted fold change for the four methods. The blue dotted line is the *x* = *y* line. In the positive, quadrant the numbers above and below the line designate the number of regressions for which the *y*-axis has a larger predicted fold change than the *x*-axis method, and vice versa. Likewise for the numbers in the negative quadrant, except here the numbers relate to absolute fold change.

**Fig S5.**
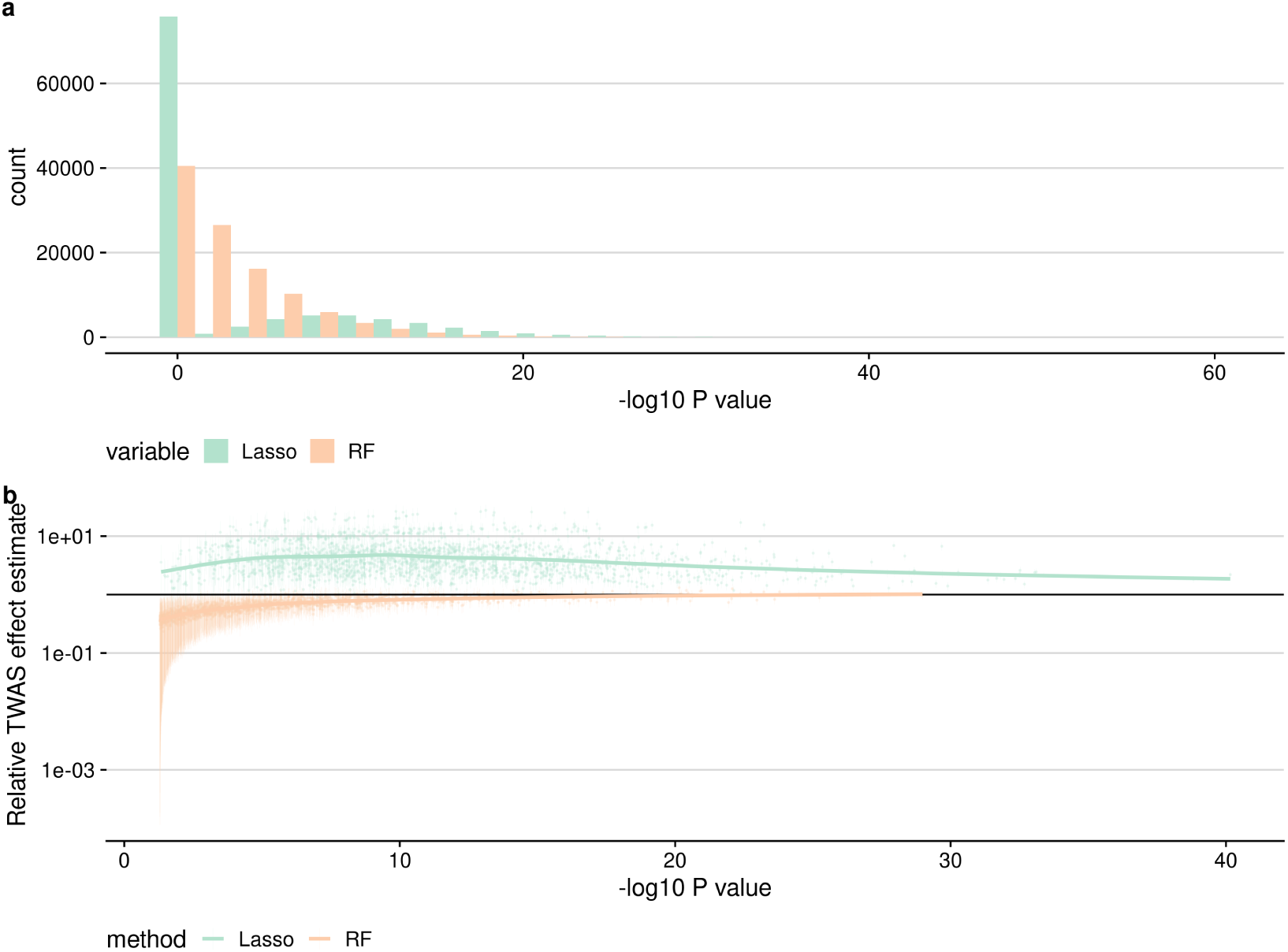
Effects of lasso regularisation on TWAS. **a** Lasso-TWAS *p*-values amongst simulations with shared eQTL/GWAS causal variants show a spike at *p*=1, and a longer tail than RF, indicating that weaker effects are missed by lasso, but that stronger effects can show greater significance compared to RF. **b** TWAS effect estimates (estimated causal effect of expression on GWAS trait) are underestimated for weak effects for RF, tending to 1 for stronger effects. For lasso, TWAS effect estimates are systematically over estimated, even for well-powered studies.

**Table S2.** Target Validation analysis of TWAS genes by method. The top 5 diseases ranked by relevance p value, and the rank of four type 1 diabetes-related terms are shown.

| Method | Disease | Rank |
| --- | --- | --- |
| Joint Lasso | hematological measurement | 1 |
| Joint Lasso | measurement | 2 |
| Joint Lasso | large intestine disease | 3 |
| Joint Lasso | intestinal disease | 4 |
| Joint Lasso | musculoskeletal system disease | 5 |
| Joint Lasso | type I diabetes mellitus | 45 |
| Joint Lasso | diabetes mellitus | 54 |
| Joint Lasso | Permanent neonatal diabetes mellitus | 1139 |
| Joint Lasso | autoimmune type 1 diabetes | 1254 |
| Lasso | type II hypersensitivity reaction disease | 1 |
| Lasso | reproductive system or breast disease | 2 |
| Lasso | carcinoma | 3 |
| Lasso | epithelial neoplasm | 4 |
| Lasso | autoimmune disease of endocrine system | 5 |
| Lasso | type I diabetes mellitus | 19 |
| Lasso | diabetes mellitus | 29 |
| Lasso | Permanent neonatal diabetes mellitus | 66 |
| Lasso | autoimmune type 1 diabetes | 731 |
| RF | autoimmune disease of endocrine system | 1 |
| RF | type I diabetes mellitus | 2 |
| RF | small intestine disease | 3 |
| RF | glucose metabolism disease | 4 |
| RF | endocrine pancreas disease | 5 |
| RF | diabetes mellitus | 6 |
| RF | autoimmune type 1 diabetes | 388 |
| RF | Permanent neonatal diabetes mellitus | 558 |
| RF-MTL | ulcerative colitis | 1 |
| RF-MTL | autoimmune disease of endocrine system | 2 |
| RF-MTL | type I diabetes mellitus | 3 |
| RF-MTL | autoimmune disease | 4 |
| RF-MTL | glucose metabolism disease | 5 |
| RF-MTL | diabetes mellitus | 12 |
| RF-MTL | Permanent neonatal diabetes mellitus | 248 |
| RF-MTL | autoimmune type 1 diabetes | 415 |

**Fig S6.**
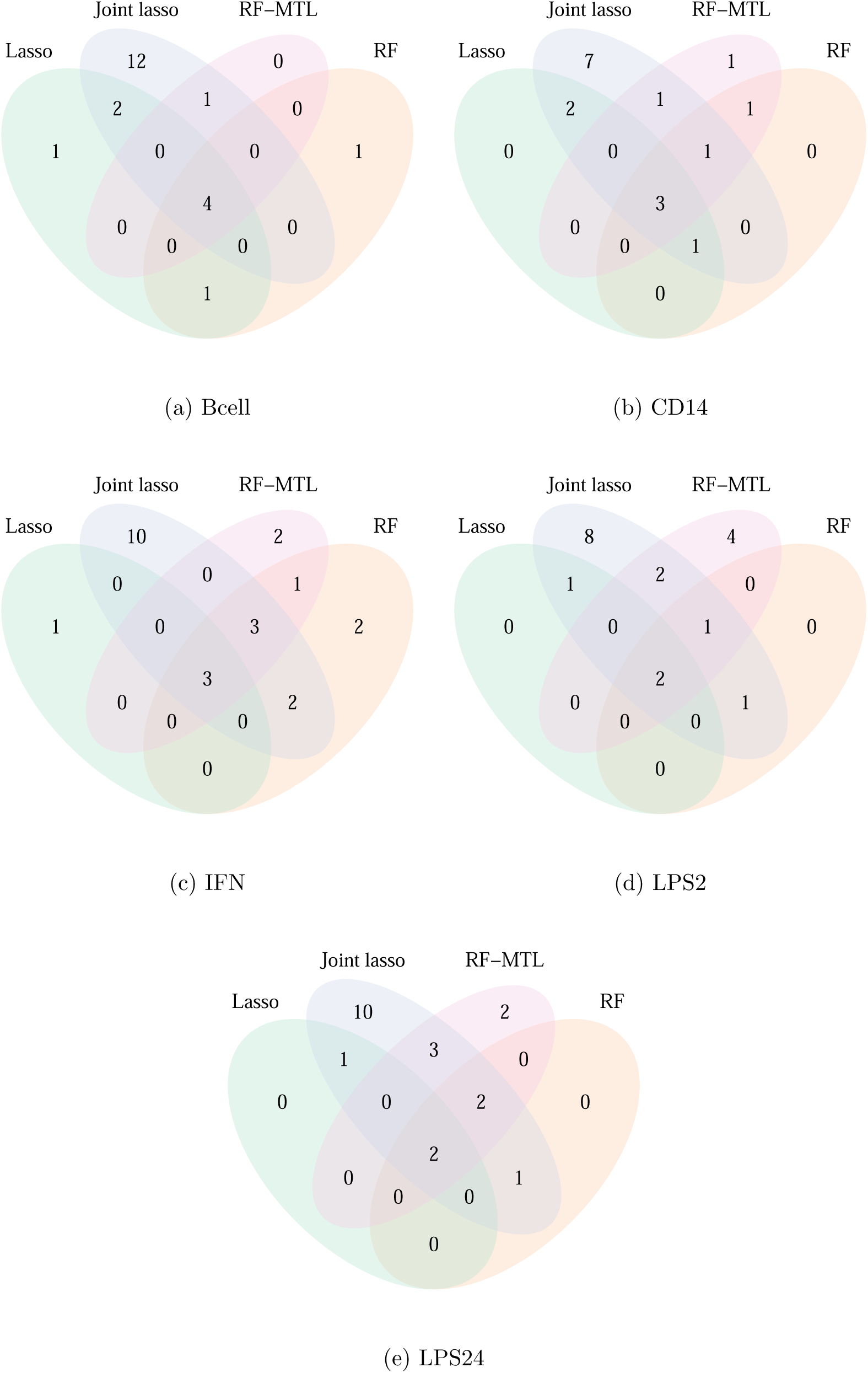
Venn diagrams showing unique SP-genes identified by the four methods for the five cells considered.

**Fig S7.**
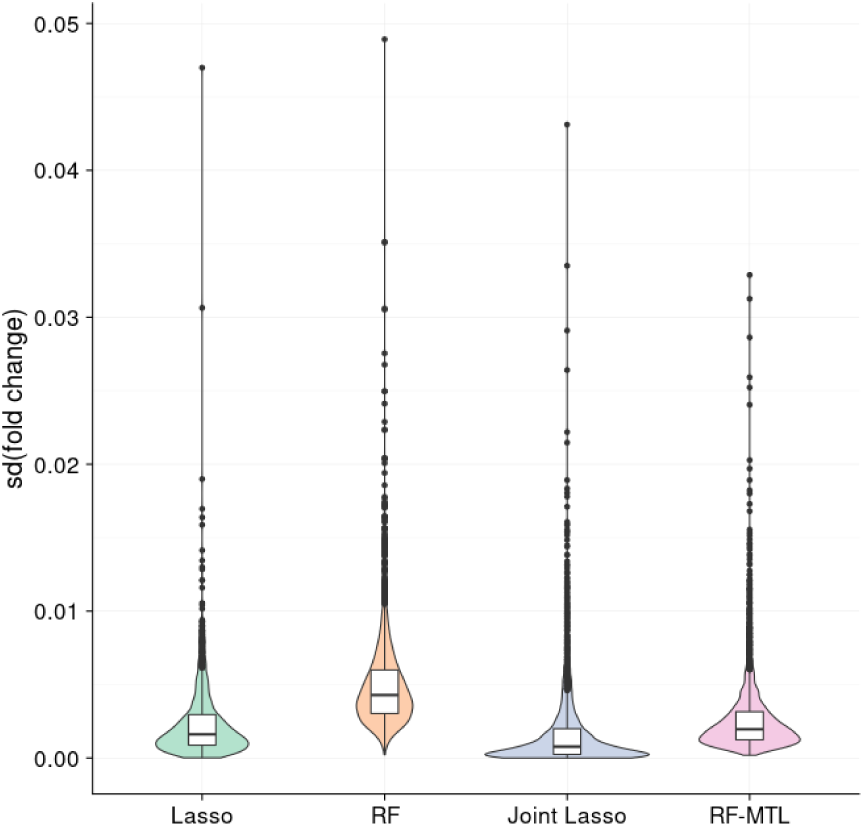
Violin plots (with inscribed boxplots) of standard deviations of predicted fold change for different cell types for each probe, per method. For each method, only probes with predictions for at least three cell types were considered.

